# Secretion and transfer of adipose lipoprotein lipase utilizes neutral sphingomyelinase 2 generated exosomes

**DOI:** 10.1101/2025.07.31.665751

**Authors:** Terri A. Pietka, Edward F. Morris, Megan Basco, Jennifer Shew, Ni-huiping Son, Zhenxiu Liu, Brandon A. Davies, Ira J. Goldberg, Clair Crewe, Nada A. Abumrad

## Abstract

Lipoprotein lipase (LPL) is critical for clearance of circulating triglycerides and for tissue fatty acid supply. LPL is primarily synthesized and secreted by adipocytes into the interstitium and must traffic from there to the abluminal/basolateral side of capillary endothelial cells. There, LPL binds glycosylphosphatidylinositol-anchored protein 1, GPIHBP1, which stabilizes the protein and facilitates its movement across the endothelial cells to the luminal side where it functions in hydrolysis of lipoprotein triglycerides. Importance of LPL traffic is supported by findings that rare mutations in GPIHBP1 cause hypertriglyceridemia. However our understanding of how LPL is secreted by adipocytes and traffics to endothelial cells is incomplete. Here we examined the possibility that secretion and traffic of adipocyte LPL might involve generation of small extracellular vesicles (sEVs/exosomes) which often mediate cell-cell communication. Proteomic analysis of sEVs secreted by adipocytes showed them enriched in LPL. To study LPL secretion and transfer we generated human derived pre-adipocytes (HPA) that stably express tagged LPL (FLAG and His epitopes). LPL pulldown and sEV isolation from HPA conditioned media documented that greater than 70% of secreted LPL is present in sEVs. The mechanism for LPL secretion in sEVs was found to involve the ESCRT-independent neutral sphingomyelinase 2 (nSMase2) pathway, as treatment with the nSMase2 inhibitor GW4869 reduced secretion by 80%. The above observations were reproduced using highly sensitive nanoparticle flow cytometry. The sEV associated LPL has lipolytic activity and it is released by heparin addition indicating it is on the sEV surface. In addition, using human derived microvascular endothelial cells with stable lentiviral expression of GPIHBP1 we show that LPL positive sEVs transfer LPL to these cells, but not to control cells without GPIHBP1. Our findings suggest that sEV formation by nSMase2 controls adipocyte LPL secretion and traffic, that sEVs protect LPL activity and facilitate LPL transfer to GPIHBP1 on endothelial cells.

## INTRODUCTION

In the circulation most fatty acids are present esterified into the triglycerides (TGs) of TG-rich lipoproteins (TRLs), chylomicrons and very low-density lipoproteins (VLDL). Lipoprotein lipase (LPL) is the key enzyme responsible for hydrolysis of TRLs^1^. Release of fatty acids from TGs by LPL is the required first step in fatty acid uptake by adipose and muscle tissues with the second step involving capture and transcytosis of fatty acids by capillary endothelial cells (ECs) for export to tissue cells^2^. LPL deficiency in humans results in severe hypertriglyceridemia (HTG), which increases risk of pancreatitis, heart disease, and Alzheimer’s^3-5^.

LPL functions in TG hydrolysis while tethered on the luminal surface of capillary ECs. The endothelium does not express LPL, which is synthesized and secreted primarily by adipocytes and myocytes^6^. Once secreted into the interstitial fluid, LPL has to transfer across capillary ECs in the tissue to localize on the EC apical/luminal surface where it can access circulating TRLs^7^. LPL localization at the apical EC surface requires its binding to the glycosylphosphatidylinositol-anchored HDL binding protein 1 (GPIHBP1) exclusively expressed by capillary ECs^7,8^. Deletion of GPIHBP1 in mice and GPIHBP1 mutations in humans markedly raise blood TGs^8,9^. Although LPL transfer from adipocytes to capillary ECs is a critical step in TG clearance, several components of this transfer remain undefined notably as related to LPL secretion and LPL traffic to bind GPIHBP1 on capillary ECs.

Adipose tissue (AT) exerts great influence on whole-body glucose and lipid metabolism through release of regulatory proteins or adipokines such as adiponectin, fatty acid binding protein and resistin. Recent evidence shows that adipocytes release many of these regulatory proteins within small extracellular vesicles, sEVs or exosomes^10,11^. The adipose derived sEVs carry metabolic proteins; lipids, metabolites and nucleic acids, etc. that can modulate function of distant recipient cells^12-14^. In turn the secretion and content of adipose sEVs are modulated by adipocyte metabolism^10,15^ consistent with the critical metabolic role of exosomal communication between adipocytes and other cells. In this study we examined whether LPL release by adipocytes might utilize sEVs. We reasoned that this could protect LPL activity and facilitate transfer of adipocyte LPL to GPIHBP1 in ECs. In this study we show that the majority of adipocyte secreted LPL is sEV-associated, and that LPL secretion in sEVs involves the neutral sphingomyelinase pathway for sEV biogenesis. The sEV-associated LPL is active and is released from sEVs by heparin. In addition, LPL on sEVs specifically binds to GPIHBP1 expressed on microvascular ECs.

## MATERIALS AND METHODS

### Cell Culture and Treatment

Human preadipocytes (HPA), isolated from adult subcutaneous adipose tissue (Sigma), were maintained in preadipocyte growth medium (Sigma) and passaged a maximum of 8 times. Human microvascular endothelial cells (MEC), isolated from adult skin (Lonza), were cultured in endothelial growth medium (EGM-2MV, Lonza) and used at fewer than five passages. Human embryonic kidney (HEK) cells (ATCC) were cultured in DMEM (Gibco) supplemented with 10% fetal bovine serum (Sigma) and 1X antibiotic-antimycotic solution (Corning). *nSMase2 Inhibition:* To inhibit nSMase2 activity and block sEV production, HPA cells were treated overnight with 10 µM GW4869 in serum-free DMEM. Condition media were collected following treatment and processed for downstream analyses.

### Lentiviral Production

To generate lentiviral particles, lentiviral constructs for tagged LPL (Addgene)^16^ and GPIHBP1^7^ were transfected into HEK cells along with 3^rd^ generation helper plasmids (Applied Biological Materials) at a 1:1 ratio using Lipofectamine 3000 (Life Technologies), following the manufacturer’s protocol. Lentiviral particles were concentrated from collected media using Lenti-X (Takara Bio) and titrated by qPCR (Applied Biological Materials).

### Overexpressing Cell Development

To generate overexpressing cells, LPL-lentivirus was transduced into HPA, AC16 and HL-1 cells using regular growth medium supplemented with 8 µg/mL polybrene. Titered lentiviral particles were added at a multiplicity of infection of 10 for all cell lines. For stable expression of GPIHBP1 in MEC, similar transduction conditions were used. The morning after transduction, cells were placed in regular growth media and cultured for 72 hours before adding 1 ug/mL puromycin for positive selection. Cells were maintained in puromycin (1 µg/mL) containing medium for all subsequent experiments to ensure continued selection.

### Extracellular Vesicle Isolation

For extracellular vesicle production, cells were incubated in either serum-free medium (HPA) or medium supplemented with reduced EV-depleted serum (AC16, HL-1) for 24–48 hours. Conditioned media were concentrated by ultrafiltration using 100 kDa cutoff filters (Millipore) to generate an EV-rich concentrate. Microvesicles were isolated by centrifugation at 18,000 × g for 45 minutes. Small extracellular vesicles (sEVs) were isolated as described below for downstream analyses.

To generate an EV-depleted conditioned media fraction, the flow-through from the 100 kDa filtration step was collected and further concentrated using 10 kDa cutoff filters (Millipore).

For heparin treatments, conditioned media were incubated with 10 units/mL heparin prior to ultrafiltration.

### EV Characterization

#### Nanotracking Analysis

Conditioned media were diluted 1:20 in 20 nm (Whatman) filtered PBS (Sigma) and analyzed for particle size and concentration using nanoparticle tracking analysis (Nanosight 300, Malvern).

#### Western Blotting

For western blotting, sEVs were precipitated using recombinant Tim4 bound to magnetic beads (MagCapture, FujiFilm). Tim4 binds to phosphatidylserine, which is enriched on the surface of sEVs. Precipitated vesicles were boiled in 1X SDS sample buffer for 3 minutes. Proteins were resolved using Bis-Tris 4–12% gradient gels (Life Technologies) and transferred to PVDF membranes (Immobilon-FL, Millipore).

For total protein normalization, membranes were stained with a total protein stain (LI-COR) prior to blocking. Membranes were blocked (1X TBS, 0.25% fish gelatin, 0.01% Na-azide, 0.05% Tween-20) and incubated overnight with primary antibodies at a concentration of 1:1000. Blots were then washed with TBST and incubated with IR-Dye secondary antibodies (1:10000) for 1 hour at room temperature and imaged ((Li-Cor Biotechnology). Primary antibodies used were anti-hLPL (R&D Systems), anti-CD81 (Santa Cruz Biotechnology), GAPDH (Cell Signal) and anti-S-tag (Life Technologies).

#### Nano Flow Cytometry

To detect LPL by flow cytometry, anti-hLPL (R&D Systems) antibody was conjugated to Alexa Fluor 647 using Mix-n-Stain chemistry (Sigma) following manufacturers protocol. For FACS analysis, 5.00×10^6^ sEV were diluted in 20-nm filtered PBS incubated overnight with 0.1 µg of anti-hLPL-AF647 and 1.6 µM CFSE (Santa Cruz Biotech). Following staining, sEVs were purified by size exclusion chromatography (CL6B300, Sigma) to separate labeled sEVs from unbound antibody and dye and eluted using 20-nm filtered PBS. Stained EVs were further diluted in filtered PBS prior to analysis using a Cytek Northern Lights instrument equipped with enhanced small particle detection (ESP) module. SSC and fluorescence gains were set to increase the resolution of EVs, and samples were run at a low flow rate using a set acquisition volume. Single stained controls were used for signal compensation. All samples were analyzed using FlowJo (v10.8.1) software.

#### EV Proteomics

Subcutaneous stromal vascular fraction (SVF) cells were isolated from 5-week-old C57BL/6 mice and differentiated into adipocytes as previously described^10^. On day 8 of differentiation, cells were washed with PBS and incubated for 24 hours in treatment media (FluoroBrite DMEM supplemented with penicillin/streptomycin, GlutaMAX, and 1% exosome-depleted FBS (System Biosciences). Conditioned media were collected, concentrated to 1 mL with 100 KD MWCO centrifugal filters (Amicon), and extracellular vesicles were isolated by size exclusion chromatography. Purified EVs were run on an SDS-PAGE gel, proteins excised and submitted to the UT Southwestern Proteomics Core^10^ for analysis.

#### LPL Activity Assay

Lipase activity of LPL-bound sEVs or conditioned media was measured using a fluorescent lipase substrate (Cell Biolabs). The substrate remains quenched until cleaved by LPL. Kinetic fluorescence measurements were performed using a BioTek Synergy II plate reader. LPL concentration was quantified using a standard curve, according to the manufacturer’s protocol.

#### LPL-GPIHBP1 Binding Assay

Control and GPIHBP1-expressing MECs were incubated with 5.00×10^6^ of LPL-sEVs for 2 hours at 4°C^17^. After incubation, cells were gently washed and lysed in cell lysis buffer (Cell Signaling Technology) supplemented with protease inhibitors (Pierce). Protein concentration was determined by BCA (Bio-Rad), and 10 µg of each lysate was separated by SDS-PAGE and probed for LPL and GPIHBP1 as described above.

### Statistical Analyses

All data are derived from a minimum of three independent experiments. Quantitative results are presented as mean values ± standard error of the mean (S.E.M.). Statistical analyses were performed using an unpaired Student’s t-test for comparisons between two groups. Statistical significance is indicated as follows: ****P ≤ 0.0001, ***P ≤ 0.001, **P ≤ 0.01, *P ≤ 0.05, and ns = not significant (P > 0.05).

## RESULTS

### LPL is present in adipocyte secreted sEVs

We recently showed that the generation of fatty acid-enriched sEVs mediates transfer of circulating fatty acids by endothelial cell CD36^18^. LPL and CD36 closely coordinate regulation of tissue fatty acid uptake and for this reason, we examined whether the sEV pathway might contribute to transfer of adipocyte released LPL. We first conducted proteomic analysis of sEVs purified from the media of cultured mouse adipocytes. The analysis verified presence of the exosome markers CD9, CD81, ALIX, SDCBP and CAV1 along with classic adipocyte markers; adiponectin, CD36, FASN, FABP, MFGE8, PLIN1 and PNPLA2 (**Table 1**). LPL was found in the sEVs and was highly enriched as compared to notable adipocyte proteins such as adiponectin and CD36. LPL enrichment in adipocyte-derived sEVs suggested that sEV generation is a potential mechanism for LPL secretion by adipocytes and LPL transfer to microvessels.

**Table 1:** Proteomics of extracellular vesicles secreted by cultured mouse derived adipocytes show sEV enrichment in LPL. Shown are proteins commonly used as sEV markers (CD9, CD81, ALIX, Syntenin-1), or that have been previously associated with sEVs (Cav-1, FABP4, MFGE8, CD36).

| Marker of | Protein Name | Relative Abundance |
| --- | --- | --- |
| *sEV* | Caveolin 1 (CAV1) | 0.939 |
| *sEV* | CD9 antigen (CD9) | 0.007 |
| *sEV* | CD81 antigen (CD81) | 0.272 |
| *sEV* | Programmed cell death 6 interacting protein (ALIX) | 0.254 |
| *sEV* | Syntenin-1 (SDCBP) | 0.108 |
| *Adipocyte* | Adiponectin (ADIPOQ) | 2.335 |
| *Adipocyte* | Fatty acid synthase (FASN) | 89.991 |
| *Adipocyte* | Heparan Sulfate Proteoglycan 2 (HSPG2) | 22.795 |
| *Adipocyte* | Lipoprotein lipase (LPL) | 13.293 |
| *Adipocyte* | Patatin-like phospholipase domain containing 2 (PNPLA2) | 0.110 |
| *Adipocyte* | Perilipin 1 (PLIN1) | 0.120 |
| *Adipocyte/sEV* | CD36 molecule (CD36) | 0.206 |
| *Adipocyte/sEV* | Fatty acid binding protein 4, adipocyte (FABP4) | 0.531 |
| *Adipocyte/sEV* | Milk fat globule-EGF factor 8 or lactadherin (MFGE8) | 5.412 |

### Human preadipocytes expressing LPL (LPL-HPA) secrete the majority of LPL in sEVs

To directly investigate LPL’s potential association with extracellular vesicles, we engineered a human pre-adipocyte (HPA) cell line that stably expresses LPL tagged with dual FLAG and His-tag epitopes (LPL-HPA). Conditioned media from LPL-HPA was confirmed to contain LPL unlike that from control HPA (**Figure 1A**). The enzyme exhibited lipase activity, which was not detected in control HPA (**Figure 1B**). Medium fractionation showed a small amount of secreted LPL (less than 30%) was associated with microvesicles, the larger particles that originate from outward budding of the plasma membrane^19^ while most medium LPL (greater than 70%) was identified in sEVs which originate from multivesicular bodies (MVB) with the sEVs released upon MVB fusion with the membrane^19^ (**Figure 1C**). Nanotracking analysis of sEVs isolated from control and LPL-HPA cells revealed that the sEVs have the expected size (86 and 96 nm, respectively) and contained the sEV marker CD81 (**Figure 1D**).

**Figure 1:**
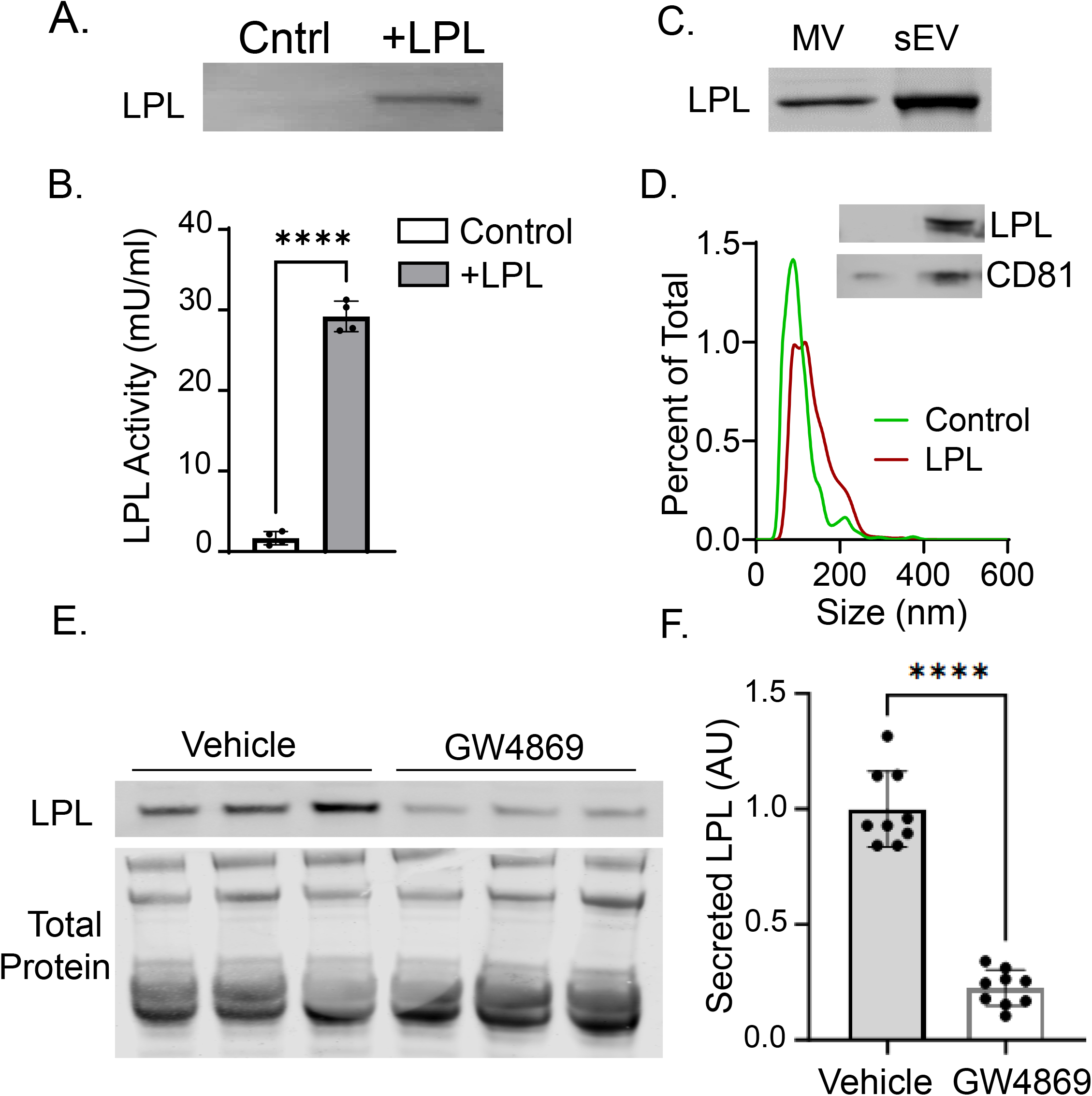
Secreted LPL predominantly associates with small extracellular vesicles (sEVs) and is dependent on neutral sphingomyelinase 2 (nSmase2) activity. **(A)** Western blot confirming the presence of LPL in conditioned media from engineered human pre-adipocyte cells expressing FLAG- and His-tagged LPL (LPL-HPA). **(B)** Assay of lipase activity demonstrates enzymatic activity in conditioned media from the LPL-HPA cells. **(C)** Fractionation of the conditioned media reveals that at least 70% of secreted LPL associates with sEVs, while ∼30% is found in microvesicles. **(D)** Nanoparticle tracking analysis (NTA) shows similar size distribution of sEVs isolated from control and LPL-HPA cells, with average diameters of 86 nm and 96 nm, respectively; CD81 is an sEV marker. **(E-F)** Western blot and densitometry analysis showing ∼80% reduction in secreted LPL levels when conditioned media is obtained from LPL-HPA cells treated with 10 μM GW4869, a specific inhibitor of nSMase2.

### Biogenesis of LPL sEVs involves the neutral sphingomyelinase 2 pathway

Exosome/sEV formation occurs through the Endosomal Sorting Complex Required for Transport (ESCRT)^20^, or through other ESCRT-independent mechanisms^21^. A critical ESCRT-independent pathway is the one controlled by neutral sphingomyelinase 2 (nSMase2), a product of the *SMPD3* gene. nSMase2 localizes to the Golgi and to the internal leaflet of plasma membrane caveolae. Active nSMase2 generates ceramide from sphingomyelins^22,18^ in the plasma membrane and in endosomes^18,21^. This facilitates caveolae endocytosis, and the budding of vesicles from endosomal membranes into the lumen of multivesicular bodies (MVBs), the luminal vesicles are then secreted as exosomes upon MVB fusion with the plasma membrane. In ECs nSMase2 controls fatty acid uptake through generation of fatty acid enriched sEVs that contain CD36 and caveolin 1 (Cav-1)^18^. Interestingly, a recent study showed that in adipose cells intracellular LPL vesicles colocalize with caveolin-1^23^.

We examined whether nSMase2 might control LPL secretion. For this, LPL-expressing HPA were treated with 10uM GW4869, a well characterized nSMase2 inhibitor that competes for nSMase2 activation by phosphatidylserine ^24,25^ and has been extensively used to inhibit nSMase2^26^. GW4869 blocks nSMase2 mediated exosome/sEVs formation. LPL-HPA conditioned media were collected, concentrated by ultrafiltration, and analyzed for LPL content via Western blot. Treatment with GW4869 as compared to vehicle resulted in 80% reduction of LPL secretion (**Figure 1E-F**).

The above findings were verified further using sensitive nanoparticle flow cytometry (nanoFACS) with sEVs isolated from control HPA and LPL-HPA cells. The sEVs were stained with <u>Carboxyfluorescein Succinimidyl Ester</u>(CFSE) and LPL was identified using a validated anti-human LPL antibody (anti-LPL-AF647). Only the sEVs secreted by LPL-HPA and not the sEVs from HPA controls were found to contain LPL (**Figure 2A-B**). The nSMase2-dependent mechanism for LPL secretion was similarly validated using nanoFACS. Treatment with GW4869 resulted in 83% reduction in the fraction of LPL-positive sEVs (**Figure 2C-D**). Together these data support the concept that most secreted LPL is associated with nSMase2 generated sEVs.

**Figure 2:**
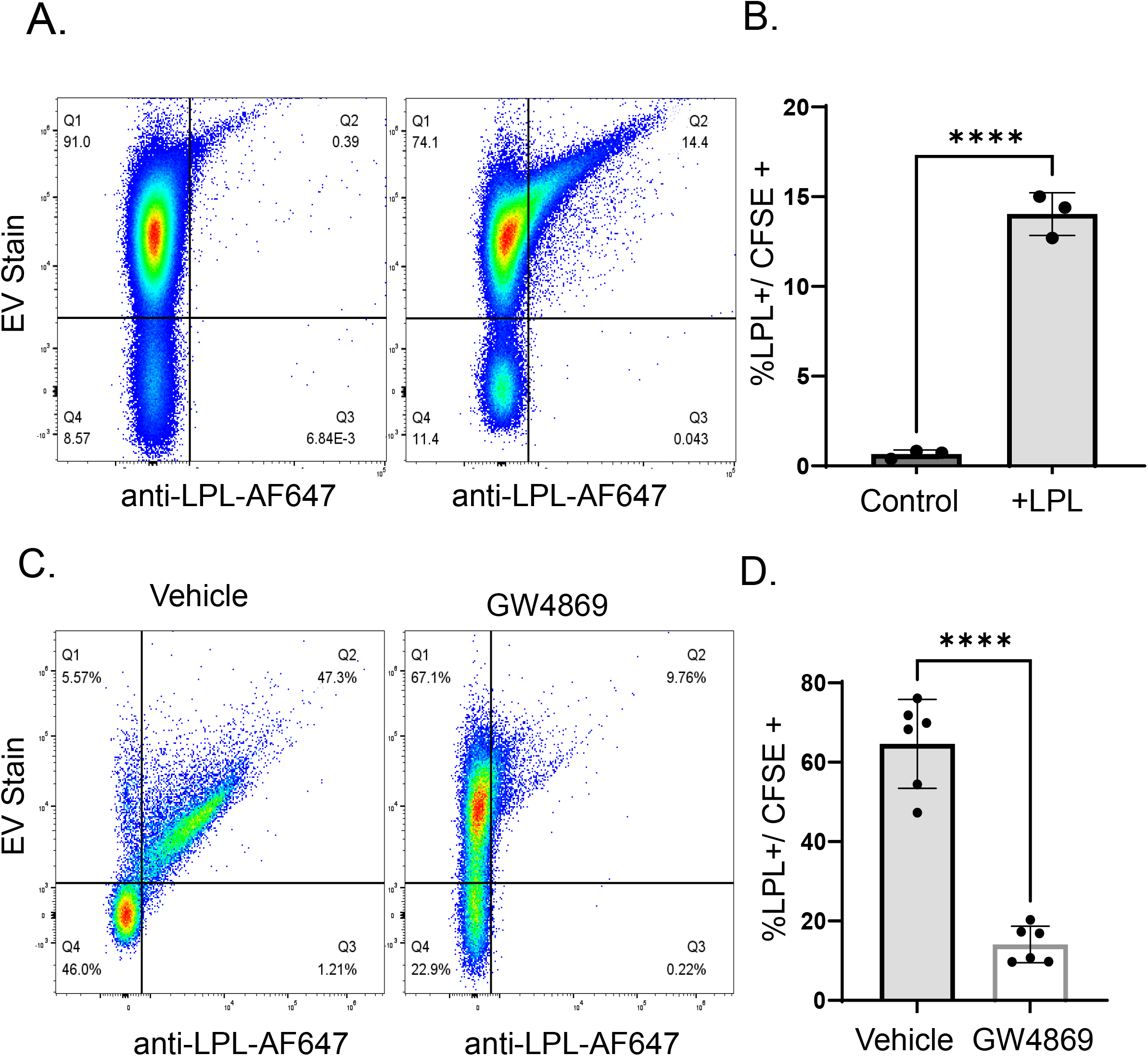
Sensitive nanoparticle flow cytometry (NanoFACS) Validates and quantifies LPL association with sEVs. **(A)** NanoFACS with sEVs preparations from control and LPL-expressing cells confirms LPL-sEV association. **(B)** NanoFACS quantification of LPL sEV association. **(C-D)** Representative nanoFACS plots of LPL+sEV signal under control and GW4869-treatment, with NanoFACS quantification showing an ∼83% reduction of LPL-positive sEVs following GW4869 treatment.

### LPL is bound to the sEV surface

To investigate the nature of LPL association with sEVs, we examined whether it is surface-bound or present within the sEV cargo. LPL contains a heparin-binding domain, and heparin treatment releases LPL from membrane heparan sulfate proteoglycans (HSPGs) and GPIHBP1^7^. We determined if sEV associated LPL can be released by heparin. Media from LPL-HPA were treated with heparin, then fractionated into microvesicles (MV), sEVs, and an sEV-depleted (-sEV) fraction (**Figure 3A**). Western blot analysis showed that heparin treatment completely dissociates LPL from MV, it reduces LPL association with sEVs and shifts the majority of LPL to the EV-depleted fraction. NanoFACS confirmed these results by demonstrating that treatment with increasing heparin concentrations caused a dose-dependent decrease in LPL positive sEVs (LPL-sEV) (Figure 3B-C). Collectively, the findings show that LPL associates with the sEV surface and can be released by heparin.

**Figure 3:**
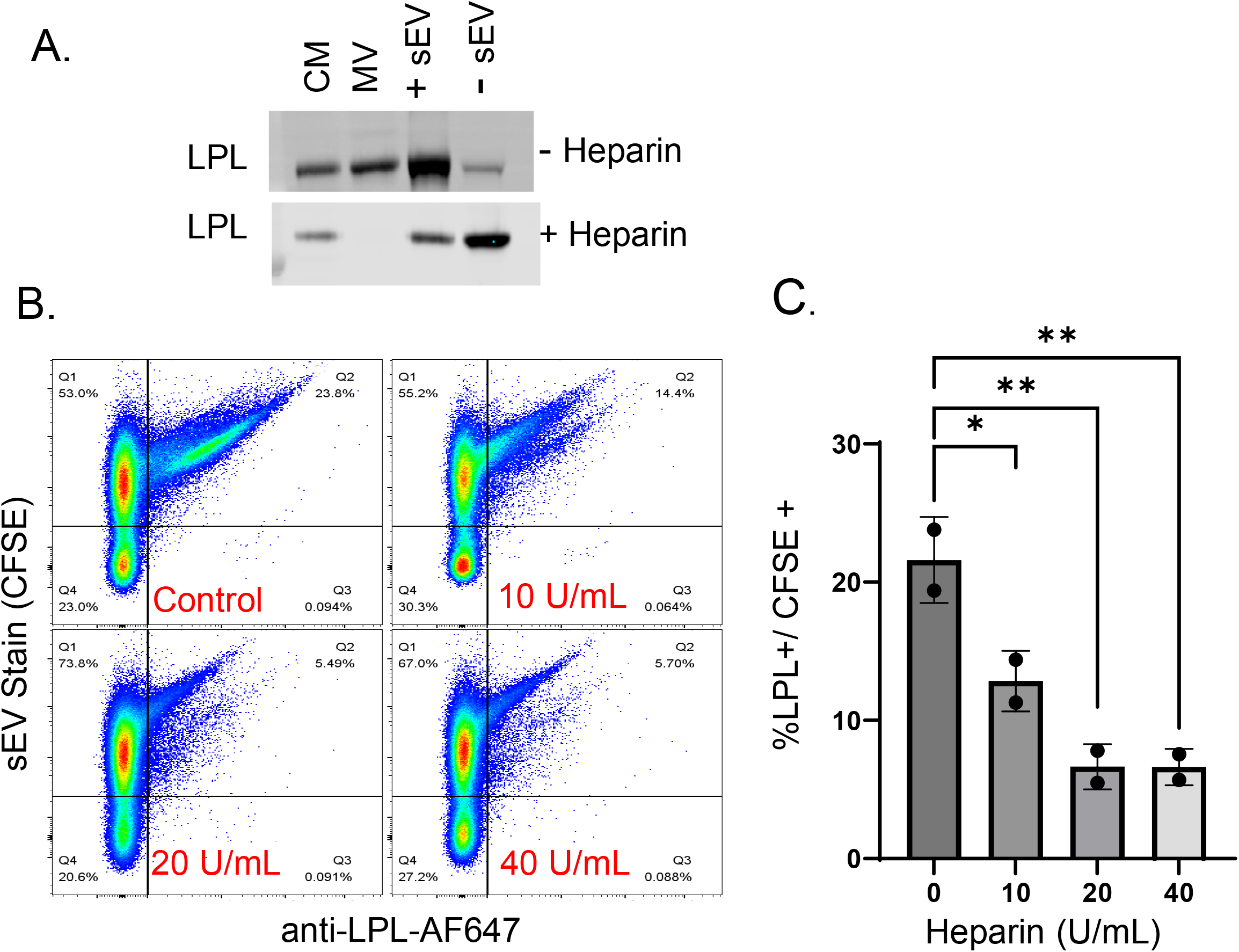
LPL associates with the sEV surface and is released by heparin treatment. **(A)** Western blot analysis of microvesicles (MV), small extracellular vesicles (sEVs), and the sEV-depleted fraction following heparin treatment of conditioned media from LPL-HPA cells. Heparin completely dissociates LPL from MV, reduces LPL levels in sEVs, and increases LPL presence in the EV-depleted fraction. **(B)** NanoFACS shows a dose-dependent reduction in LPL-positive sEVs at increasing concentrations of heparin, and the quantification of the progressive loss of LPL signal from sEVs **(C)**.

### sEV-associated LPL exhibits lipolytic activity

LPL activity is known to be unstable in cell culture media so its presence on the surface of sEVs raises the question of whether sEV associated LPL is enzymatically active. Using a fluorescent LPL activity assay, we measured LPL activity in size-exclusion purified sEVs from control and LPL-HPA cells. No activity was detected in control sEVs, whereas activity was measured in sEVs from LPL-HPA (**Figure 4A-B**). We also assessed LPL activity in conditioned media following its fractionation into sEV-enriched and sEV-depleted fractions using ultrafiltration. No activity was detected in the sEV depleted fraction (**Figure 4C-D**). The above data show that sEV associated LPL is active and they are consistent with the sEVs serving as functional carriers of the secreted active LPL.

**Figure 4:**
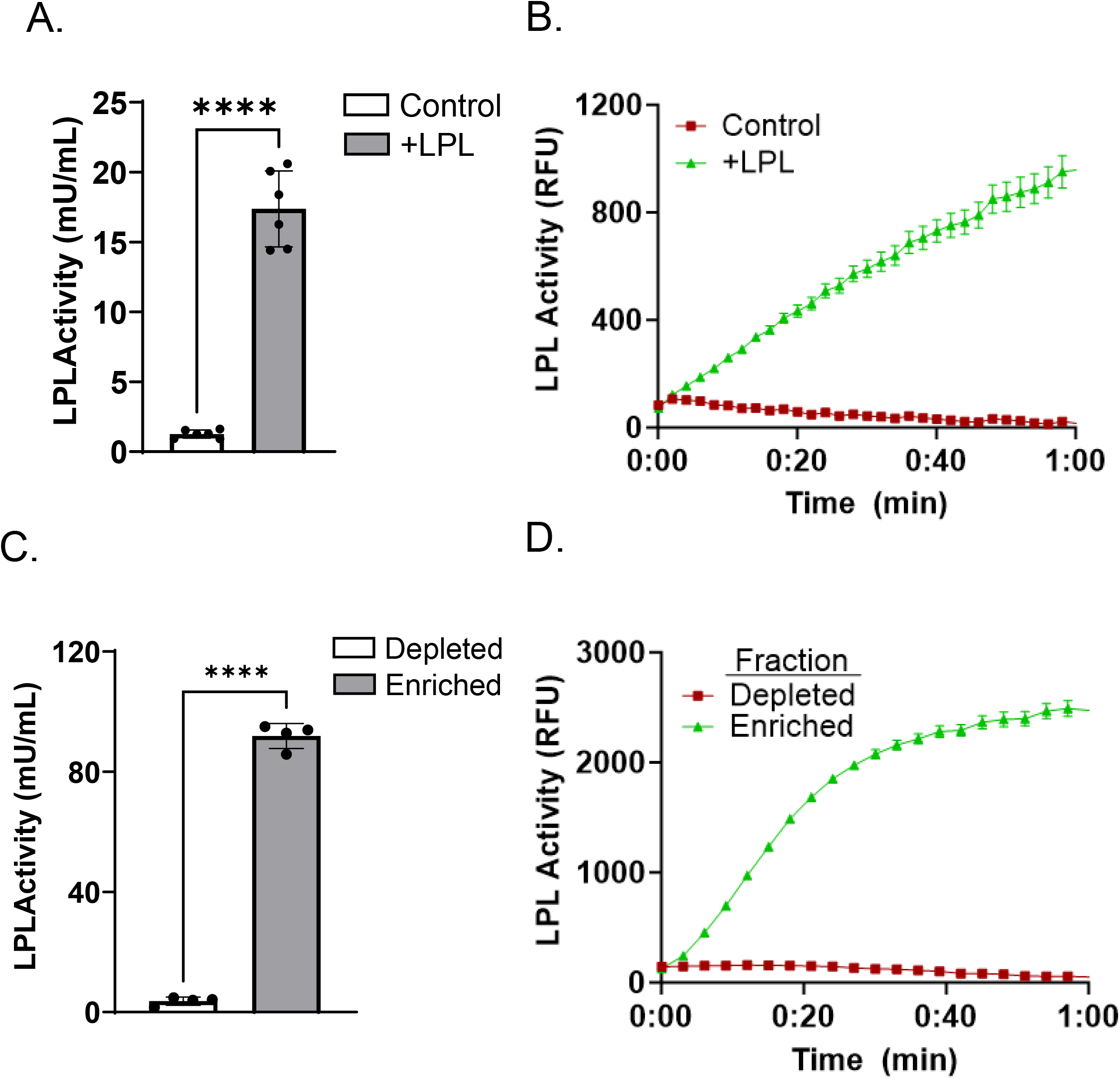
The sEV-associated LPL has enzymatic activity. **(A)** Fluorescent lipase activity assay shows no detectable activity in size-exclusion purified sEVs from control cells. **(B)** sEVs from LPL-HPA cells exhibit measurable lipase activity, indicating enzymatically active LPL on the sEV surface. **(C)** Conditioned media from LPL-HPA cells were fractionated by ultrafiltration into sEV-enriched and sEV-depleted fractions. **(D)** LPL activity was detected exclusively in the sEV-enriched fraction, and no activity is observed in the sEV-depleted fraction.

### The sEVs mediate transfer of LPL to endothelial cell GPIHBP1

Transfer of LPL from the abluminal EC side to its functional site on the luminal surface of ECs involves GPIHBP1. This transfer is critical to ensure efficient LPL lipolysis of circulating TG-rich lipoproteins. We examined whether sEVs can deliver LPL to GPIHBP1. Microvascular cells rapidly lose GPIHBP1 expression in culture, so we generated microvascular ECs with stable GPIHBP1 expression (GPIHBP1-ECs) using a lentiviral system. Control and GPIHBP1-expressing ECs were incubated with LPL-containing sEVs. The ECs were then carefully washed, and levels of LPL and GPIHBP1 assessed by western blot. We observed low LPL binding to control ECs, whereas GPIHBP1-expressing cells bound substantially more LPL (**Figure 5A-5B**). Since GPIHBP1 promotes LPL stabilization and transport to the apical surface, these findings suggest that the sEVs facilitate transfer of active LPL to GPIHBP1 on microvascular ECs.

**Figure 5:**
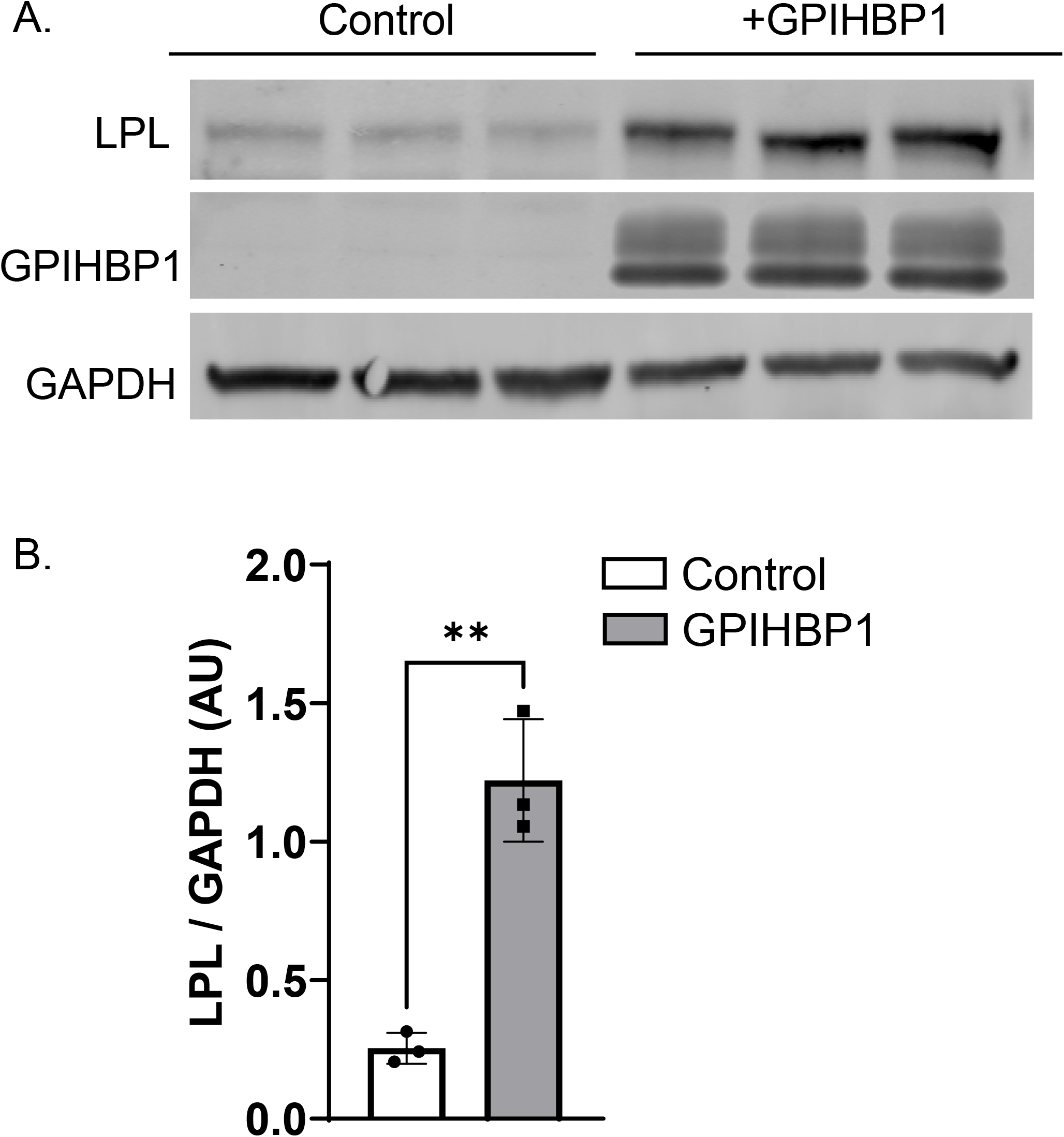
sEVs mediate transfer of LPL to endothelial cells via GPIHBP1. **(A)** Western blot analysis of LPL levels in control and GPIHBP1-expressing microvascular endothelial cells (GPIHBP1-ECs) following incubation with LPL-containing sEVs. **(B)** Quantification of LPL binding shows significantly greater LPL association with GPIHBP1-ECs compared to control ECs, consistent with sEV delivery of LPL to GPIHBP1 on endothelial cells.

## DISCUSSION

LPL modulates the metabolism of both TGs and fatty acids, it acts to clear circulating TGs preventing the deleterious effects of persistent hypertriglyceridemia and together with CD36, LPL coordinates FA supply to tissues. Adipocytes are a major provider of LPL which must transfer to microvascular ECs to function in TG hydrolysis. It is well established that LPL binding to GPIHBP1 and to heparan sulfate proteoglycans (HSPG) plays an important role in its transfer to ECs^7,8,27^. This study adds new information to the transfer mechanism for adipose LPL. First, it documents that secretion of LPL by adipocytes and LPL transfer from adipocytes to microvascular endothelial cells involve biogenesis of sEVs, with the LPL-sEV pathway responsible for transport of a substantial fraction of secreted LPL. Second, LPL is released from the sEVs by heparin indicating its presence on the sEV surface where it is likely bound to heparan sulfate proteoglycans, present on sEVs^28,29^. Third, the sEV-associated LPL is lipolytically active, suggesting its sEV transport protects its activity. Fourth, the LPL-sEVs can deliver LPL to microvascular ECs expressing GPIHBP1, but not to control ECs with no GPIHBP1 expression.

Adipose tissue (AT) maintains homeostasis of energy metabolism through its function to store or release fatty acids in response to changes in the nutritional milieu. In addition, AT secretes metabolic modulators that influence other organs, notably the nano-sized exosomes or sEVs. Adipose sEVs are now recognized as potent metabolic modulators and have been documented to play an important role in cell-cell communication ^30,31^ and in the regulation of systemic metabolism ^12,14^ by acting through transfer of bioactive cargo that includes proteins, lipids, metabolites, RNA, DNA, etc^12,32^. The sEVs cargo typically includes bioactive components that influence the recipient ECs reinforcing importance of AT-EC crosstalk^12^ and of the metabolic role of the adipose secretome. Our current findings on the role of adipose released sEVs in LPL secretion and traffic are consistent with the metabolic regulatory role of AT.

Our findings suggest that the sEV pathway could be exploited to help explain unidentified causes of hypertriglyceridemia, and subsequently for therapeutic approaches as sEVs offer the possibility of specific cell targeting^33^ with protected LPL activity. Obesity associates with alterations of sEV production and sEV content^34-36^ and also with abnormal LPL regulation, the increase in LPL activity observed in response to feeding fails to occur in individuals with obesity^37^ and could not be explained by altered LPL mRNA. The absent increase could potentially involve a defect in sEV generation and LPL secretion.

The control of LPL-sEV biogenesis by nSMase2 is of interest. Although nSMase2 has been primarily studied in pathological conditions, we had recently identified its role in EC fatty acid uptake^18^. This together with the current findings support a key contribution of the enzyme to regulation of fatty acid metabolism. nSMase2 localizes to plasma membrane caveolae and to the Golgi apparatus. Its role in sEV formation is mediated by generation of ceramide in the membrane and in endosomes, which facilitates both endocytosis and the sorting of endocytosed vesicles for secretion^18,21,25^. Our findings suggest that reduced adipose nSmase2 activity could elevate circulating TGs by impairing LPL secretion and traffic. Conversely, reduced EC nSMase2 activity may elevate circulating fatty acids since EC fatty acid transfer plays a limiting role in tissue fatty acid uptake^38^. Understanding how adipose nSMase2 is regulated may provide insight into the factors that determine normal versus abnormal TG clearance. For example, in ECs, nSMase2 is activated by fatty acids^18^ and by high glucose^39^.

It is important to note that our studies with secretion and transfer of adipose LPL might not apply to muscle and heart cells which also synthesize LPL. Muscle LPL activity correlates with LPL mRNA levels^40^, whereas activity in AT is exclusive of mRNA or protein levels^41,42^. Locally produced angiopoietin-like protein 4 (ANGPTL4) inactivates adipose LPL and its expression is downregulated postprandially when storage of TG-derived fatty acids would be expected to increase ^43^. In contrast, activity in muscle appears modulated not only by LPL production but by LPL inhibition by the ANGPTL3/8/aApoA5 complex, primarily derived from the liver^44,45^. Determining whether these factors differentially affect sEV associated LPL versus LPL secreted from myocytes is a goal of future studies.

In summary we identified sEVs generated through nSMase2 as the main pathway for transfer of active adipose LPL to ECs in microvessels. This provides a framework for studying pathway regulation under different metabolic conditions. LPL deficiency in mice or humans causes severe hypertriglyceridemia^1,40^. Similarly, GPIHBP1 deletion in mice and a rare GPIHBP1 mutation in humans markedly raise blood triglyceride levels^46^. In humans, mutations in LPL or GPIHBP1 are rare and cannot explain the hypertriglyceridemia commonly observed in individuals with obesity. Possibly, obesity-associated hypertriglyceridemia results from a defect in regulation of the adipocyte sEV pathway for LPL secretion, consistent with the altered sEV production and content observed in obesity ^34-36^. Improving adipocyte sEV secretion could become a therapy to enhance adipose tissue lipolysis, reduce circulating triglyceride levels, and improve insulin sensitivity by promoting more efficient lipid handling.

## ACKNOWLEDGEMENTS

The authors thank the Nutrition Obesity Research Center Cellular and Molecular Biology Core (NORC CMBC) for their technical support with these studies.

## CONTRIBUTION STATEMENT

TAP, IJG and NAA designed the studies, analyzed the data and prepared the manuscript. TAP, EFM, MB, JS, NS, XL performed the LPL experiments, CC performed the proteomics analysis on EVs. BSJD provided consultation and reagents for GPIHBP1 experiments and reviewed the manuscript. NAA is the guarantor of this work.

